# On the role of melanistic coloration on thermoregulation in the crepuscular gecko *Eublepharis macularius*

**DOI:** 10.1101/2023.05.18.541382

**Authors:** Brandon T. Hastings, Anastasiya Melnyk, Mehrdad Ghyabi, Emma White, Frederico M. Barroso, Miguel A Carretero, David Lattanzi, Julien Claude, Ylenia Chiari

## Abstract

Body coloration in ectotherms serves multiple biological functions, including avoiding predators, communicating with conspecific individuals, and involvement in thermoregulation. As ectotherms rely on environmental sources of heat to regulate their internal body temperature, stable melanistic body coloration or color change can be used to increase or decrease heat absorption and heat exchange with the environment. While the function of melanistic coloration for thermoregulation has been found to increase solar radiation absorption for heating in many diurnal ectotherms, research on crepuscular and nocturnal ectotherms is lacking. Since crepuscular and nocturnal ectotherms generally absorb heat from the substrate, coloration is likely under different selective pressures than in diurnal ectotherms. We tested if the proportion of dorsal melanistic body coloration is related to differences in body temperature heating and cooling rates in the crepuscular gecko *Eublepharis macularius* and whether changes in environmental temperature trigger color changes in this species. Temperature measurements of the geckos and of the environment were taken using infrared thermography and temperature loggers. Color data were obtained using objective photography and a newly developed custom software package. We found that body temperature reflected substrate temperatures, and that the proportion of melanistic coloration has no influence on heating or cooling rates or on color changes. These findings suggest that, in *E. macularius*, melanistic coloration may not be used for thermoregulation. Future research should further test the function of melanistic coloration in other crepuscular and nocturnal vertebrates to understand the evolution of melanistic pattern in animals active in low light conditions.

## INTRODUCTION

Optimal body temperature is essential for animals for mating, foraging, predator escape, and other biological functions (Dunham et al., 1989; Seebacher & Franklin, 2005). In the face of global climate change and frequent temperature fluctuations caused by climate change (Rummukainen, 2012; Vasseur et al., 2014), organisms will be exposed to recurrent suboptimal temperatures. Thus, examining the performance of organisms under suboptimal temperatures is essential to predict how their biology will be affected (Angilletta, 2009; Deutsch et al., 2008; M. Kearney et al., 2009; Mader et al., 2022; Pérez i de Lanuza et al., 2018). Ectotherms, such as non-avian reptiles, generally do not have a built-in physiological mechanism for internal heat production; hence, the maintenance of their body temperature, and thus of the temperature-dependent basic life functions, relies on a thermal exchange with their environment (M. Kearney et al., 2009; Kiefer et al., 2007).

Many ectothermic vertebrates have evolved a wide range of behavioral, physiological, and morphological characteristics that aid in maintaining close to functionally optimal body temperatures. Although microhabitat changes and posturing are the most common thermoregulatory behaviors observed in ectotherm vertebrates (Aubret & Shine, 2010 and references within; Bauwens et al., 1996; Kearney & Predavec, 2000), skin coloration may also influence thermoregulation. In areas with low mean annual solar radiation, ectotherm vertebrates may have higher concentrations of dermal melanin caused by the concentration of melanophores to increase heat absorption, the so-called Thermal Melanism Hypothesis (Clusella Trullas et al., 2007; Clusella-Trullas et al., 2008; Forsman, 1995; Gibson & Falls, 1979; Walton & Bennett, 1993). Vertebrate ectotherms living in areas with low levels of solar radiation – i.e., areas further from the equator – are more uniformly dark in color due to the melanophores distributed relatively equally throughout the dermis - to create an overall lower skin reflectance and higher rate of heat absorption. Ectotherms occurring in areas of higher solar radiation, however, are less constrained in increasing heat absorption from solar radiation and can exhibit clusters within their dermis that have a higher concentration of melanophores relative to other areas, causing for example a spotted pattern (Martínez-Freiría et al., 2020; Moreno Azócar et al., 2015; Szydłowski et al., 2020). Similarly, ectotherms that are mostly active during hours of low solar radiation – i.e., crepuscular and nocturnal species may rely on other strategies to absorb heat (Garrick, 2008) and melanism in these species may be used for other functions, including for cooling rather than heating (Geen & Johnston, 2014). Together with stable melanistic coloration, melanosomes may move within the dermal melanophores, producing a skin darkening when moving towards the surface of the skin, or skin lightening when moving away from it, a phenomenon called rapid physiological color change (Sherbrooke & Frost, 1989). Rapid physiological color change may be used for thermoregulation by darkening or lightening the skin surface of ectotherms and increasing or decreasing the level of solar radiation absorption and/or body heat exchange with the environment, thus altering the internal temperature of the organism (Lowe & Norris, 1956; Sherbrooke et al., 1994; Sherbrooke & Frost, 1989; Smith, Cadena, Endler, Kearney, et al., 2016; Smith, Cadena, Endler, Porter, et al., 2016). Although thermal melanism has been studied within and among species in ectothermic vertebrates, thermal melanism has been measured as overall darker or lighter skin colors, without inferring how the proportion of melanistic patterns may influence the rates of heating and cooling and the internal body temperature of the organism (though see Forsman, 1995 for snakes). Furthermore, to our knowledge, physiological color change in response to temperature changes influenced by the proportion of melanistic pattern (i.e., melanistic and non-melanistic coloration becoming lighter or darker at different rates) has not been examined.

The leopard gecko, *Eublepharis macularius,* a species with widespread melanistic pattern (Glimm et al., 2021), is characterized by melanistic spots (at the adult stage) or stripes (at the juvenile stage) on a lighter, often yellow, background of base dermal coloration. It is also commonly bred for pet-trade resulting in a variety of melanistic patterns of different types and quantity (Glimm et al., 2021; Kiskowski et al., 2019; Szydłowski et al., 2020) that is ideal for testing the influence of melanistic proportion on biological processes. To our knowledge, no study has assessed and quantified rapid physiological color change in this species. *Eublepharis macularius*, a species that naturally occurs in Pakistan, India, Iran, and surrounding regions (Agarwal et al., 2022), is a crepuscular lizard primarily active in low light conditions. As such, we expect that heat absorption in this organism does not strongly depend on solar radiation – as it is active during times of low solar radiation – and that dorsal melanistic proportion would not affect heating rates in this species. On the other hand, melanistic proportion may have an influence on increasing cooling rates during especially hot days, while the organism is hiding. We expect *E. macularius* to be thigmothermic – absorbing heat from the direct contact with warm substrates while hiding during the day or moving on surfaces that have absorbed heat during the day at night.

In this work, we used multispectral objective photography (Troscianko & Stevens, 2015) and a newly developed data extraction pipeline to extract the melanistic coloration from the background animal coloration on live and free to move *E. macularius*. Although other algorithms and pipelines are available to extract color data, including segmenting color patterns from background colors (see Glimm et al., 2021, Abramjan et al., 2020, and Troscianko & Stevens, 2015), our approach allowed for extracting the same type of color data over time – i.e., every 30 min at which the images of the gecko were taken - on the same freely moving individual allowing coloration and color pattern to be compared over time. In this study, we tested if: 1. individuals with a greater proportion of stable dorsal melanistic pattern across the body experience increase heating or cooling rates than individuals with lower proportion; 2. exposing individuals to lower suboptimal temperatures for this species elicits a physiological color change in the skin of the geckos; 3. individuals with a greater baseline proportion of melanistic coloration experience a lower amount of overall physiological color change (background light color and melanistic color). Experimental results on this model are expected to contribute to elucidate the function of melanistic coloration and physiological color change in this species, and further our understanding on the role of melanistic coloration on thermoregulation vs. crypsis in crepuscular and nocturnal species.

## MATERIALS AND METHODS

All capture, handling, and experimental protocols were approved by George Mason University IACUC committee (Permit number 1718778). Experiments were carried out to minimize stress and disturbance to the animals and in accordance with relevant guidelines and regulations.

### Study subject and captivity conditions

All geckos were housed in the same room at George Mason University, with one gecko per terrarium (61x30x20 cm) and exposed to a 12-12hr day-night cycle. The housing containers for *E. macularius* were plastic boxes with newspaper bedding. The room temperature was kept stable between 25-28°C, with the average temperature generally being close to 25.3°C +/-0.2°C, and each terrarium contained moist and dry hides and a heat pad for thermoregulation. Humidity (41+/-10%) and temperature in the room were checked daily using a digital thermometer/hygrometer. The health condition of each individual was checked daily by visual inspection, and no gecko was tested during the shedding process. Geckos were fed three times a week with a combination of crickets and mealworms dusted with calcium and vitamin powder. Feeding was withheld 3 days prior to testing in order to avoid interference in digestion due to exposure of animals to low temperature and to also avoid confounding effects due to potential food digestion. Feeding was resumed after testing. Drinking water was always available except during testing. Testing was carried out on 12 adult geckos (n=12, 7 males, 5 females). None of the females used in this experiment were gravid, and neither any of the tested individuals had been engaging in mating or any social interaction as each individual was housed and tested in isolation from the others.

### Experimental Setup

Experiments were conducted in a temperature controlled room of 4x2 m at George Mason University. This room has a 1x3 m open area in the middle, where the 51x25x30 cm glass terrarium used as a testing enclosure was placed on a rubber mat on the floor. The walls and floor of the terrarium were covered with white Teflon (SS Shovan) to remove any potential effect of background coloration, and the lid was removed for the entirety of the experiment to facilitate obtaining the data. The Teflon on the bottom of the terrarium consisted of three layers, while the one on the side was a single layer. Black electrical tape (3M) was used to adhere the bottom sheet of Teflon to the side sheets to prevent geckos from hiding underneath of the Teflon. The use of black electrical tape was chosen following guidelines on thermal camera calibration (F. Barroso, pers. comm.). A clean cardboard egg carton of 15x15 cm was placed in the middle of the terrarium as a hiding spot for the gecko, because reptiles are known to thermoregulate behaviorally by seeking shelter in suboptimal temperatures (Aubret & Shine, 2010; M. Kearney et al., 2009; Woods et al., 2015). Except for the cardboard egg and three iButtons (DS1921G Thermochron, Maxim Integrated Products, precision = 0.5°C) to record the temperature (see below), nothing else was placed in the terrarium and the gecko was free to move and use the entire space available in the terrarium. A heating pad was placed underneath one side 15cm from the end of the terrarium (to have a warm spot throughout the experiments). A broad-spectrum UV-VIS light (Zoo Med PowerSun H.I.D Metal Halide UVB Lamp, 6500K, 70W, 95CRI) was placed 160cm above the center of the testing terrarium to ensure photographs were taken under proper lighting (Troscianko & Stevens, 2015), as melanin has strong absorption in the UV-Vis spectrum (McNamara et al., 2021). This light was turned on 45 minutes prior to the start of each experiment and remained on for the length of the experiment (Supplementary Materials Fig. S1).

A thermometer/hygrometer (ThermoPro TP50, precision=0.1°C, 1% humidity) was placed on the floor adjacent to the terrarium to measure the temperature and humidity of the room before and during the experiments. To monitor the temperature throughout the experiments and to ensure that room and terrarium temperatures were similar across experiments, a total of four iButtons were placed in the terrarium and in the temperature controlled room. Specifically, one iButton was placed on the floor outside of the middle section of the terrarium 20 cm away. Of the three iButton placed in the terrarium, two were placed on the opposite ends (one on top of the heating pad and the second on the opposite end of the terrarium) and one under the cardboard egg carton. The iButtons were programmed to start collecting temperatures data 30 minutes prior to the start of the experiment and continued to collect temperature data every five minutes for the entire duration of the experiment. Following each experiment, iButtons were sanitized with isopropyl alcohol and the terrarium was cleaned with soap and hot water to remove any potential scent or residue left from the previously tested individual. The top layer of the Teflon at the bottom of the terrarium was replaced after each experiment, while the Teflon sheets on the sides were sanitized with isopropyl alcohol after each experiment. A new cardboard egg carton was used for each tested individual.

### Native environmental temperatures for *E. macularius*

To estimate the native environmental temperatures for *E. macularius* and compare them with the temperatures tested in this study, occurrences for this species were downloaded from the Global Biodiversity Information Facility (GBIF, October 2022; www.gbif.org) and imported into Rstudio (V4.1.2, R Core Team 2021) using the “occ_download_get” function from the *rgbif* package (Chamberlain et al., 2022). Species occurrences for *E. macularius* were filtered for species’ scientific name mismatches as well as NA values for latitude, longitude, species’ scientific names, and country codes. Species occurrences were also cleaned and cross-checked for coordinate validity using the “clean_coordinates’’ function from the package, *Coordinate Cleaner* (Zizka et al., 2019). Species occurrences which resulted in at least one flagged test labeled as, “FALSE,” were removed from the dataset. Microclimate temperatures were extracted using the global model from the *NicheMapR* package (M. R. Kearney & Porter, 2017). Microclimate temperatures represent temperatures at 3 cm above ground with full sun (no shade) using 23 native coordinates for leopard geckos from GBIF. The model was run over 365 days for 10 years. Microclimatic temperatures for each day were taken as the average of each 60-min time interval across the day.

### Experimental temperature ranges

The range of experimental temperatures used in this experiment was 15-25°C (Table 1). For the purposes of analysis, time periods in the experiment were separated into five “blocks” to measure differences in variables between different phases of the experiment (Table 1). Briefly, blocks 1 and 5 correspond to the beginning and end of the temperature experiment, block 2 corresponds to the temperature going down, block 4 corresponds to the temperature going up back to 25°C, and block 3 corresponds to the lower temperature used in this study (Table 1). The highest temperature was chosen as 25°C because this is the overall most frequent temperature at which these geckos are exposed in their housing environment. 15°C was used as the lower temperature to resemble natural low temperature experienced by this species in its natural habitat (Fig. 1), without eliciting hibernation (Khan, 2009). Experiments focused on studying the effects of lower temperatures as *E. macularius* is a crepuscular species that is less active during the warmer parts of the day.

**Figure 1.**
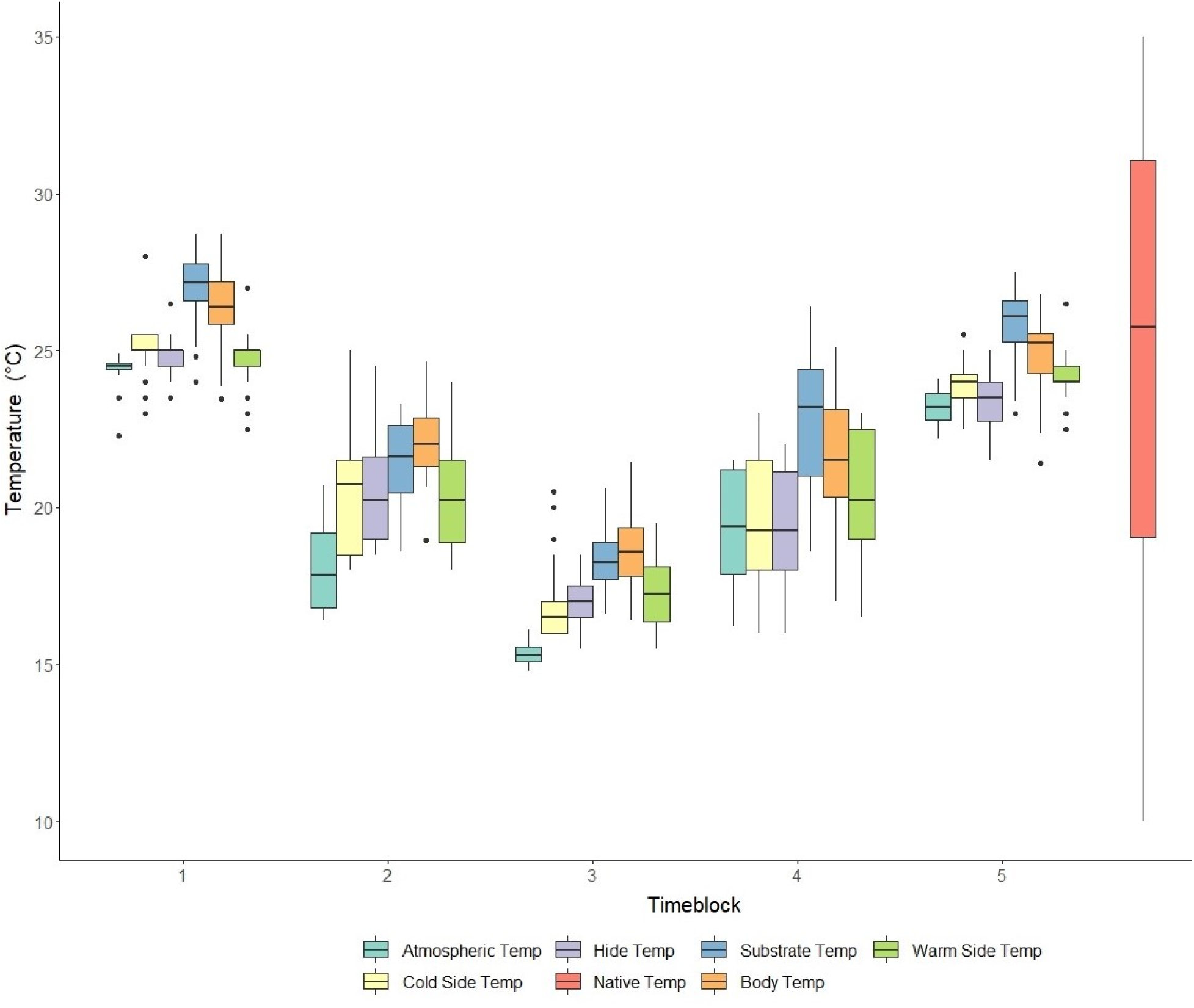
Box plots of the individual body temperatures, temperatures within terrarium and atmospheric temperatures for each block. Temperature values for each individual were plotted to visualize temperature variations for within the terrarium, atmospheric temperatures, and body temperatures in comparison to the native temperature range of *E. macularius* for each block. Blocks are as in Table 1. Body and substrate temperatures are based on IR (infrared) images. Cold side, hide, and warm side of the terrarium and atmospheric temperatures are based on temperatures recorded by dataloggers. Native temperatures are based on Global Biodiversity Information Facility. The colors used for each block correspond to the different temperatures are indicated in the figure legend.

**Table 1:** Selected experimental temperatures at different time points in the experiment. Blocks were used for the statistical analyses. Blocks were selected to minimize temperature variance within blocks and maximize variance across them. For actual experimental minimum and maximum temperatures for each block, see Fig. 1.

| **Block** | **Experiment time**<br>**(min)** | **Terrarium**<br>**temperature (°C)** | **Terrarium cold**<br>**end (°C)** | **Terrarium warm**<br>**end (°C)** |
| --- | --- | --- | --- | --- |
| 1 | 0-120 min | 25 | 25 | 25 |
| 2 | 121- 180 min | 25 decreasing to 15 | 25 decreasing to 15 | 25 decreasing to 18 |
| 3 | 181-300 min | 15 | 15 | 18 |
| 4 | 301-360 min | 15 increasing to 25 | 15 increasing to 25 | 18 increasing to 25 |
| 5 | 361-420 min | 25 | 25 | 25 |

To set up the maximum and minimum temperatures for this experiment, the temperature control for the experimental room was set at 25°C and then lowered by setting the room temperature control at 15°C when needed for the experiments (Table 1); temperature lowering between 25 and 15°C (or vice versa) took one hour. To bring the temperature back to 25°C from 15°C a space heater was placed 1 m from the warm end of the terrarium and turned on after the temperature control of the room was adjusted to 25°C (block 4, Table 1). At the beginning of each experiment, we used the room thermometer (ThermoPro) to confirm that the room temperature was at 25°C. The temperature of the terrarium was confirmed for each end of it by pointing an infrared thermometer (Etekcity Corporation) held 30 cm from the surface pointing perpendicular towards the bottom of the terrarium. Temperature checks of the room and terrarium were repeated every 30 minutes during the experiment using the in-room thermometer and the infrared thermometer, respectively. Temperature readings from iButtons were used to confirm atmospheric temperature readings after each experiment.

### Data collection

Geckos were tested in a random order. Individuals were weighed to the nearest 0.01g. using a digital scale before the start of the experiment and snout vent length (SVL) was measured to the nearest 1mm. Only one gecko per day was tested within the same time frame for 7 hours, starting at 11:00am each day. Although this species is crepuscular, experiments were carried out during the day to replicate the conditions of the housing room, where geckos are exposed to light conditions during the day. At the end of the experiment, each gecko was returned to its housing terrarium. Geckos were visually monitored after the experiments to check for any health concern. No geckos had any issues during or after the experiments.

To extract temperature data from multiple body parts of the gecko, as different body parts may have different temperatures, a CAT S62 smartphone Pro camera (Caterpillar Inc., resolution=12MP, emissivity=0.95) was used to take an infrared (IR) image (Barroso et al., 2016). The camera was held approximately 30 cm directly above the individual in order to maintain the same effective pixel size (i.e., the actual area each pixel represents in the photographed subject) across IR images, regardless of the body size of the animal or its position in the terrarium. To standardize IR images and determine reflective temperatures, an 8x8 cm square piece of wrinkled aluminum foil was placed next to the gecko in the terrarium when capturing the image of a gecko each time an IR image was taken, following Barroso et al. (2016). The square piece of aluminum foil in each IR image was used to extract average reflective temperature from IR images only and was not used in color analysis. Average reflective temperature is required in each IR image to standardize temperatures for the gecko and the terrarium. After taking the IR image of the gecko, the relative humidity and temperature of the temperature-controlled room were recorded.

After taking the IR image of the gecko, to obtain the color data for each gecko, visible images were taken using a full spectrum converted Canon 1300D with a Kolari Vision UV/IR cut filter (410-700nm transmission). Images were taken approximately 40cm directly above the gecko to ensure a good resolution across images and with a grayscale standard built from Teflon following the methods of Abramjan et al. (2020) in the frame of the image. Visible images were obtained only for the dorsal part of the geckos, as melanistic patterns are generally absent from the ventral side of the animals and as such the ventral side was not relevant to the study questions (Glimm et al., 2021). Two people (AM and EW) took the IR images making sure to standardize the method between them and another person (BH) always took the visible images. If the gecko was under the cardboard hide, the hide was lifted prior to taking the images and replaced afterwards. The time at which the temperature of the room was changed, the thermostat/thermometer readings, as well as the times at which visible and thermal photos were taken were manually recorded.

### Data extraction

Temperature measurements from the IR images were extracted in FLIR Tools (Teledyne FLIR 2022) from five body parts of each gecko (head – base of parietal scales, left knee, left foot, central dorsum, eyes – right and left, snout, and tail–- above the cloaca), as body temperature is known to vary across the body (Barroso et al., 2016). Temperature data averaged from both eyes were used as the internal body temperature for the analyses, as suggested for lizards by Barroso et al. (2016). Overall, the average of the temperature between two eyes and other body regions showed strong correlation, with a Spearman correlation coefficient above 0.98 (p<0.001, Supplementary Material Table S1). Furthermore, the average temperature from both eyes was found to be highly correlated with the temperature of the snout (r_s_ =0.96) which has also been suggested to be a good proxy for internal body temperature (Tabh et al., 2021). For each IR image, relative humidity, atmospheric temperature, distance (0 m), and emissivity (1) were first entered into FLIR Thermal Studio following Barroso et al. (2016) and Barroso (pers. comm.) to calibrate the temperature readings of the thermal camera. Reflective temperature was obtained as the average reflective temperature of the aluminum foil standard. This value was extracted from the IR image of the aluminum foil by overlaying a box entirely over the aluminum foil using the *Rectangle* function in FLIR Tools. After calibrating and entering the average reflective temperature in FLIR, distance and emissivity were re-entered as 0.3 m and 0.96, respectively (Barroso et al., 2016). The substrate temperature of the terrarium was also measured by using the *Rectangle* function to overlay a small box over the black electrical tape at the bottom of the terrarium for each image taken, as the electrical tape more accurately reflects the temperature of the terrarium (F. Barroso, pers. comm.).

Digital images obtained with the Canon camera were processed using custom image processing software written in the Python programming language; color space conversions and luminance calculations were done using the OpenCV package in Python (Bradski, 2008). Luminance was used as a measure color change as this is the most important color component that affects solar radiation absorption (Smith, Cadena, Endler, Kearney, et al., 2016). First, images were normalized for potential changes in lighting conditions across images of the same individual for the different blocks by converting each image to Hue -Lightness -Saturation (HLS) color space and using the grayscale reference in the first image of that individual as the baseline. The Lightness parameter (HLS) of following images were then standardized so that the greyscale reference matched that of the first image of that individual. This standardization step therefore allows comparisons among images taken for the same individual, but not across individual. All images were then converted back to the RGB color space. Next, because the limbs were sometimes obscured from the camera view due to the posture of the gecko in the image, images were cropped to only include the head, trunk, and tail of each individual. For image color segmentation – to segment the studied areas into color regions-, hierarchical k-means clustering was run on each image (the specific parameters used can be found in the available codes hosted on GitHub), as manual object segmentation can be inaccurate and time consuming, especially when dealing with a large number of images. In this process every pixel is assumed to be a datapoint in the RGB color space, then pixels are grouped into a predefined number of clusters based on their distances to each other in the three-dimensional RGB color space. Because the result of this method is sensitive to its initialization, cluster centers were initiated following a K-means++ algorithm to account for this (Arthur & Vassilvitskii, 2007). While the K-means algorithm is computationally inexpensive and fast, there were some limitations caused by lighting conditions with a high incidence of shadows. It was common for coloration in areas of discoloration due to shadowing to be incorrectly assigned to a cluster, resulting in a misrepresentation of the pattern for that image. Because of the unsupervised nature of k-means, there is no way to correct for this error once the segmentation step is started. To account for this effect, a visual confirmation step by the user was implemented before segmentation to ensure that the color clusters would accurately represent the pattern to be segmented. For the present study, k=2 was used for clustering. While this is conservative, it avoided overestimation of the amount of melanistic coloration (Supplementary Materials Fig. S2).

The results of K-means clustering for visible images was a Boolean mask representing stable melanistic and non-melanistic coloration of the entire body for each image. Melanistic proportion was calculated as the area of stable melanistic coloration relative to the entire dorsal area (head, trunk, and tail), averaged across all images (15) of a single individual. The average standard deviation of melanistic proportion across all 12 geckos - calculated as average of the standard deviation for each gecko based on the 15 images taken for each individual’s melanistic proportion - was +/-3.5%. Variation in melanistic proportion could be attributed to variation in lighting conditions, different positioning of the animal across the 15 images, imaging cropping, and the fact that K-means is an iterative, unsupervised algorithm (Lloyd, 1982). The Boolean mask was applied to the color corrected image to extract the mean luminance value for melanistic and non-melanistic coloration and the proportion of the coloration that was either melanistic or non-melanistic relative to the total coloration of the gecko. Luminance values were derived from a weighted calculation of RGB color channels (specific information are included in the data extraction pipeline on GitHub). All Python codes used to extract color data are available on GitHub. The full dataset will be publicly available on Dryad *after manuscript acceptance*.

### Statistical analyses

Pearson’s correlation coefficients were used to evaluate the relationship between the gecko body temperature and the terrarium substrate temperature, both calculated from the same IR image, or between the gecko body temperature and the average of the three datalogger temperatures placed inside the terrarium, or between the gecko body temperature and the atmospheric temperature as estimated on the data logger outside the terrarium. To confirm the accuracy of IR substrate readings, a Pearson’s correlation test was also run on the average data logger temperatures from within the terrarium and the substrate temperature taken with IR imaging. Because one temperature value from the data loggers had a reading of zero, that value was removed from the analyses. Previous studies have found that humidity may influence body temperature (Galliard et al., 2021); as such, we also tested the potential correlation between humidity and body temperature using a Pearson’s correlation test.

To test for overall differences in body temperature throughout the experiment, we performed paired t-tests or Wilcoxon-tests (based on normality of the data) between the average body temperatures of each gecko in each block. The rate at which geckos heat and cool down may be influenced by body weight or by the animal size; as such, a linear model was fit between body weight and heating and cooling rates separately and the analyses were then repeated for SVL. Furthermore, as different sexes may respond differently to heating and cooling, we also used a linear model to test the influence of sex effects on heating and cooling rates separately. Heating and cooling rates were calculated taking the change in body temperature between blocks 1 and 3 (cooling) and blocks 3 and 5 (heating).

To ensure that there was no correlation between the proportion of dorsal melanism (taken for each gecko as the average log of melanistic proportion over the 15 images) and snout-vent length (SVL) or sex, a linear model was fit between melanistic proportion and SVL with sex as a factor. Melanistic proportion was transformed to log of melanistic proportion to obtain normality of the data. To investigate the influence of melanistic proportion on heating and cooling rates, we ran a linear model based on data obtained on all 12 geckos between the log of the average melanistic proportion taken for each gecko and heating/cooling rates as defined above.

Paired T-tests or Wilcoxon-tests (depending on normality of the data) were used to evaluate any changes in luminance values (physiological color change) between blocks. To assess the influence of body temperature variation and log of the average melanistic proportion on physiological color changes (luminance) on the entire dorsal area (head, trunk, and tail) of the gecko, we used linear models with the luminance change between two blocks as the dependent variable with body temperature change between two blocks or log of averaged melanistic proportion of each individual as the independent variable. We also ran more general models where these two independent variables (melanistic proportion and body temperature) were jointly tested with their interaction. As the results do not vary between the simpler and more complex model, we only report the results of the complex model to also assess the influence of the interaction between melanistic proportion and body temperature. Analyses were run independently for changes between blocks 1 and 3 (cooling), 3 and 5 (heating), and 1 and 5 (initial and final body temperatures). Luminance for melanistic and non-melanistic parts of the body of each gecko was tested in separate models. Any statistically significant linear model results were investigated further with diagnostic tests, specifically the Cook’s distance to evaluate how much leverage each individual exerted on the model. All statistical analyses were run in R (V4.1.2, R Core Team 2021).

## RESULTS

For each of the 12 tested geckos, 15 IR and 15 visible images were used for the analyses, with the exception of one IR image each for two geckos being of too low quality for data extraction, giving 178 IR data points and 180 visible data points (15 x 12 individuals). Similarly, 179 temperature measurements were used for each temperature type (IR images or average of the three dataloggers inside the terrarium) for the analyses as well. All temperatures taken from the experiments were evaluated against the geckos’ native temperature range based on WorldClim data (Fig. 1). Experimental temperatures stayed within the quartiles of native temperature as planned by the experimental design, with the starting and ending temperature of 25°C corresponding to the median of native temperatures (Fig. 1). As a consequence, blocks 1 and 5 are closest in temperature to the interquartile range of the native temperature (Fig. 1), while block 3 had the greatest temperature difference from the median native temperature. Specifically, in block 3 all the experimental temperatures have a median temperature lower than the lower quartile of the native temperatures (warm side of the terrarium=17.25°C, cold side=16.5°C, hide spot=17°C, native lowest quartile=19.04°C; p<0.001 based on Wilcox tests for all data logger temperatures compared to native low quartile of native temperature), supporting that the temperatures we selected as suboptimal in this study are in fact suboptimal for this species in its native environment.

The geckos’ body temperature was strongly correlated with the terrarium substrate temperature (r^2^= 0.97, p=0.33), both obtained from IR imaging. Correlation between the geckos’ body temperature and the atmospheric or terrarium temperatures based on data loggers were also strong (r^2^= 0.92 or 0.93 depending on the comparison, p<0.001 for all correlations). Although, correlation between body temperature and each of the three dataloggers placed in the terrarium (cold side, warm, and hide) and between body temperature and atmospheric have identical r^2^, the median atmospheric temperature is generally lower than the body temperature and the terrarium temperature based on data loggers (Fig. 1). Finally, correlation between the average temperature of all the three dataloggers within the terrarium and the substrate temperature estimated from the IR images were also high (r^2^= 0.85, p=0.40), although the temperature estimated by the IR images was relatively higher for each block than the one based on the datalogger (Fig. 1). We also found that relative humidity was not correlated with body temperature (r^2^=-0.23, p=0.82). To notice that while the median gecko temperature at the beginning and end of the experiment is lower than the median substrate temperature estimated by IR imaging, as the temperature of the experiment decreases (block 2-3), the median body temperature is higher than the median substrate temperature and closer to the lower quartile of the native temperatures (Fig. 1).

Significant differences in body temperature were detected during cooling (blocks 1 and 3, t-test t=41.2, p=2.08x10^-13^), heating (blocks 3 and 5, Wilcox test V=0, p= 0.0005), and between beginning and end of the experiment (blocks 1 and 5, Wilcox test V=78, p=0.0005) (Table 2). The heating and cooling rates were found to be independent of body weight (r^2^=-0.02 p=0.39 for heating and r^2^=0.19 p=0.09 for cooling), SVL (r^2^=-0.09 p=0.76 for heating and r^2^=0.06 p=0.22 for cooling), or sex (r^2^=-0.05 p=0.50 for heating and r^2^=-0.07 p=0.60 for cooling) (Table 3).

**Table 2.** Differences in body temperature (a), and luminance for the melanistic (b) and non- melanistic (c) part of the body during cooling, heating, and between the beginning and end of the experiments. Cooling occurs between blocks 1-3, heating during blocks 3-5, and beginning and end of the experiment between blocks 1-5. More information about blocks specific temperatures can be found in Table 1. Depending on the data distribution, paired t-tests or Wilcox test were run; as such we report corresponding t-value for t-tests and V-value for Wilcox test. Significant p-values (<0.05) are indicated in bold.

| Block comparisons | df | t-value | V | p-value |
| --- | --- | --- | --- | --- |
| a) Body temperature |  |  |  |  |
| 1-3 | 11 | 41.232 | NA | **2.076x10<sup>-13</sup>** |
| 3-5 | NA | NA | 0 | **0.00055** |
| 1-5 | NA | NA | 78 | **0.00055** |
| b) Luminance for melanistic body areas |  |  |  |  |
| 1-3 | 11 | 1,5257 | NA | 0.16 |
| 3-5 | 11 | -1.0559 | NA | 0.31 |
| 1-5 | 11 | 0.84798 | NA | 0.41 |
| c) Luminance for non-melanistic body areas |  |  |  |  |
| 1-3 | 11 | 1.0359 | NA | 0.32 |
| 3-5 | 11 | -0.34277 | NA | 0.74 |
| 1-5 | 11 | 0.94218 | NA | 0.37 |

**Table 3.** Influence of snout-vent length (SVL), sex, and body weight on melanistic proportion (a), heating rates (b), and cooling rates (c). In (b) and (c), we used the simplest models testing one variable at the time. Melanistic proportion was calculated as the area of stable melanistic coloration relative to the entire dorsal area (head, trunk, and tail) across all the 15 images of a single individual. Melanistic proportion was transformed as log of melanistic proportion for the analyses. Cooling rates correspond to the difference in body temperature between blocks 1 and 3, while heating rates correspond to blocks 3 and 5.

| Factors | Estimate | Std Err | t-value | p-value |
| --- | --- | --- | --- | --- |
| a) Melanistic proportion |  |  |  |  |
| SVL | -0.9906 | 1.4138 | -0.701 | 0.50 |
| Sex | -16.584 | 22.826 | -0.727 | 0.49 |
| Sex* SVL | 1.3674 | 1.9087 | 0.716 | 0.49 |
| b) Heating rates |  |  |  |  |
| Weight | -0.0670 | 0.0753 | 0.890 | 0.39 |
| SVL | -0.1315 | 0.4189 | -0.314 | 0.76 |
| Sex | -0.3546 | 0.5087 | -0.697 | 0.50 |
| Melanistic proportion | -0.0451 | 0.1542 | -0.292 | 0.78 |
| c) Cooling rates |  |  |  |  |
| Weight | 0.0980 | 0.0517 | 1.898 | 0.09 |
| SVL | 0.3897 | 0.2997 | 1.300 | 0.22 |
| Sex | 0.2126 | 0.3954 | 0.538 | 0.60 |
| Melanistic proportion | -0.1458 | 0.1099 | -1.327 | 0.21 |

Of the 12 geckos tested, nine had a melanistic proportion between 0% and 11% across the entire dorsal part of the body, while the other three had a melanistic proportion of 16%, 22%, and 41%. We found no influence of SVL (r^2^=-0.27, t=-0.70, p=0.50), sex (r^2^=-0.27, t=-0.73, p=0.49), or their interaction (t=0.71, p=0.49) on melanistic proportion (Table 3). Based on linear models, the melanistic proportion had no influence on heating rates (blocks 3 and 5, r^2^=-0.09, t=-0.292, p=0.78) or cooling rates (blocks 1 and 3, r^2^= 0.06, t=-1.327, p=0.21) (Table 3, Fig. 2).

**Figure 2.**
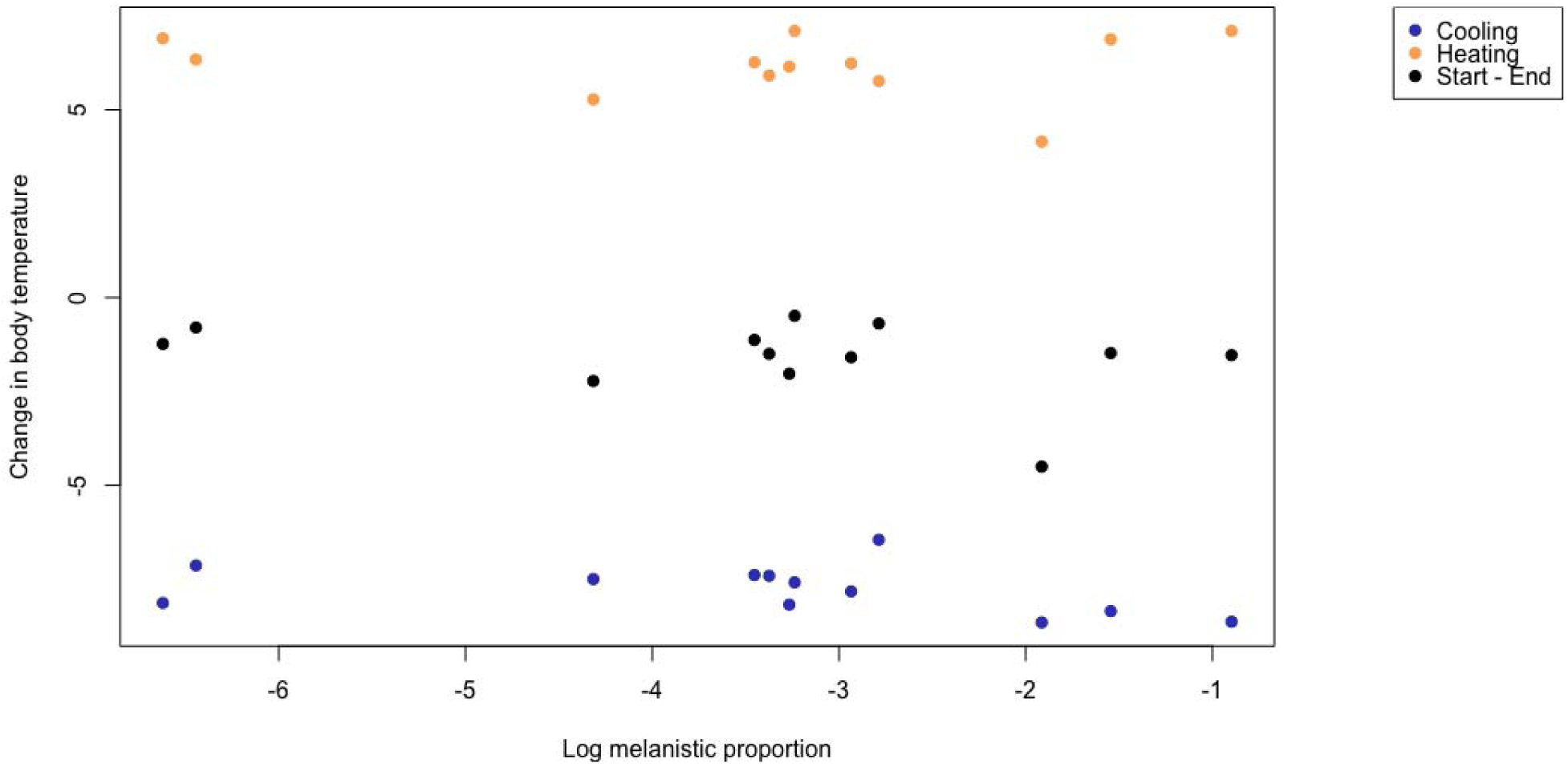
Influence of melanistic proportion on the change in body temperature. Logarithm of the average stable melanistic proportion of each individual was used for this plot. Heating rates were calculated by taking the average body temperature difference between each block for blocks 5 and 3 per individual. Cooling rates were calculated using the same methods for blocks 1 and 3. Each dot represents an individual. Blue dots refer to the cooling phase (blocks 1-3), orange dots to the heating phase (blocks 3-5), and black dots refer to the start and end of the experiment (blocks 1-5). Changes in body temperature are similar within each group (cooling, heating, start-end) independently of the melanistic proportion of each individual.

Using t-tests to evaluate differences in luminance between blocks, we found no significant differences between blocks 1 and 3 (cooling), blocks 3 and 5 (heating), and beginning and end of the experiment (blocks 1 and 5) for the melanistic and non-melanistic areas of the body (Table 2b and c for p-values). Linear models ran to test the influence of the proportion of stable melanistic coloration and body temperature changes on changes in luminance for the melanistic and non-melanistic dorsal areas of the body indicate that during heating (blocks 3-5), the melanistic proportion has an influence on change in luminance for the non-melanistic area of the body (r^2^=0.45, t=-2.861, p=0.02, Table 4). Specifically, individuals with a greater melanistic proportion had higher luminance (luminance increases, thus the animal becomes lighter in color) in the non-melanistic areas of the body, while the melanistic part of the body did not experience any change in luminance (Figs 3A and 4A). Furthermore, we found an interaction between melanistic proportion and body temperature on changes in luminance during heating (blocks 3 and 5) for the non-melanistic areas of the body (r^2^=0.45, t=2.877, p=0.02) (Table 4, Figs 3A and 4A). Diagnostic tests revealed that one individual exerted significant leverage on the results of the model and influenced the significant results (Cook’s distance > 1). Melanistic proportion and body temperature had no influence on changes in luminance during cooling (blocks 1-3) or beginning and end of the experiment (blocks 1-5) for the non-melanistic area or the melanistic area of the body and for heating and the melanistic part of the body (Table 4).

**Figure 3.**
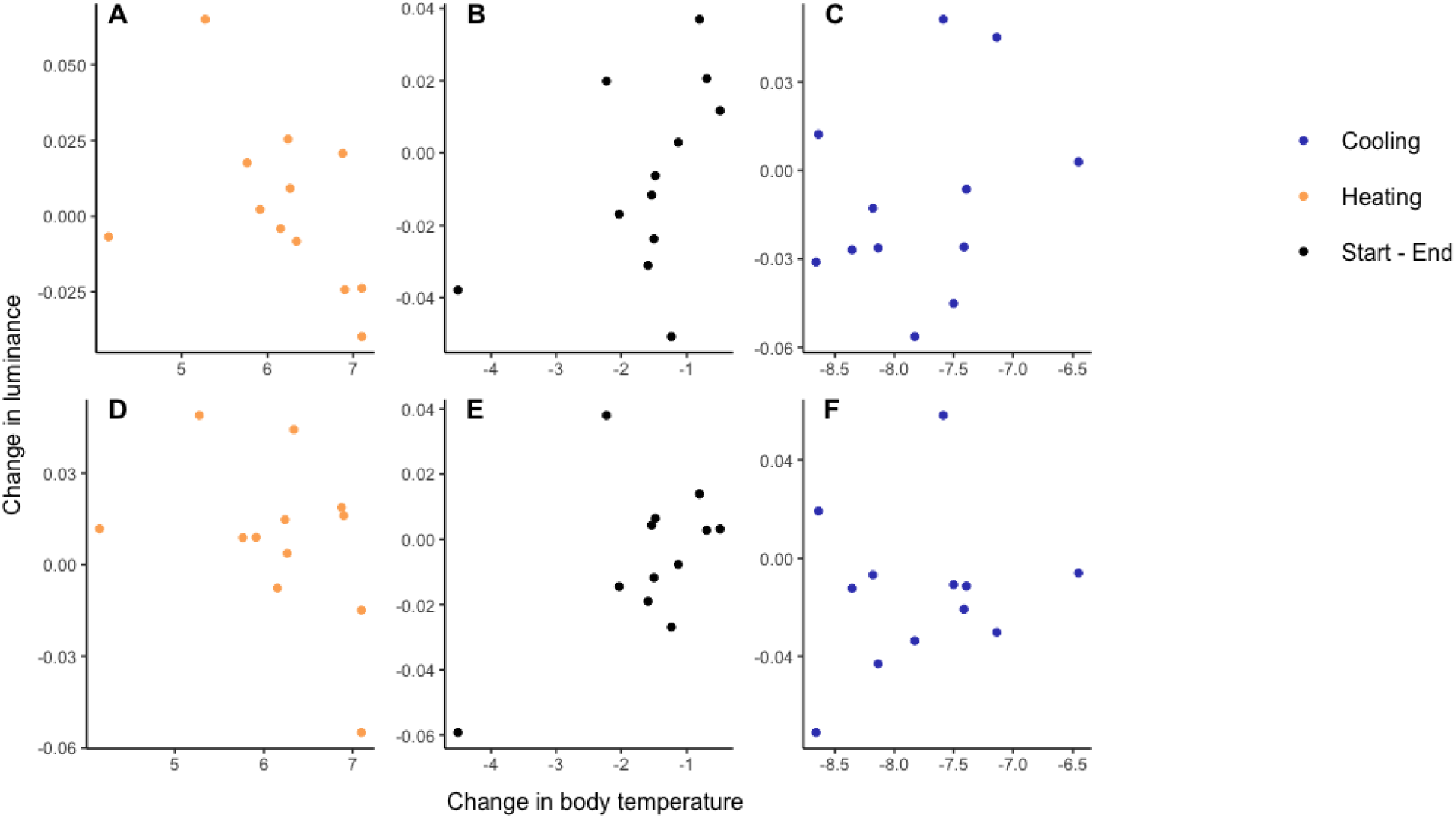
Change in average luminance for non-melanistic (A, B, C) and melanistic (D, E, F) areas of the body against change in body temperature. Average luminance was calculated by weighted RGB values taken from images. Heating rates were calculated by taking the average body temperature difference between each block for blocks 5 and 3 per individual. Cooling rates were calculated using the same methods for blocks 1 and 3. Each dot represents an individual. Blue dots refer to the cooling phase (blocks 1-3, plots C and F), orange dots to the heating phase (blocks 3-5, plots A and D), and black dots refer to the start and end of the experiment (blocks 1- 5, plots B and E). Solid line indicates a significant p-value via linear models.

**Figure 4.**
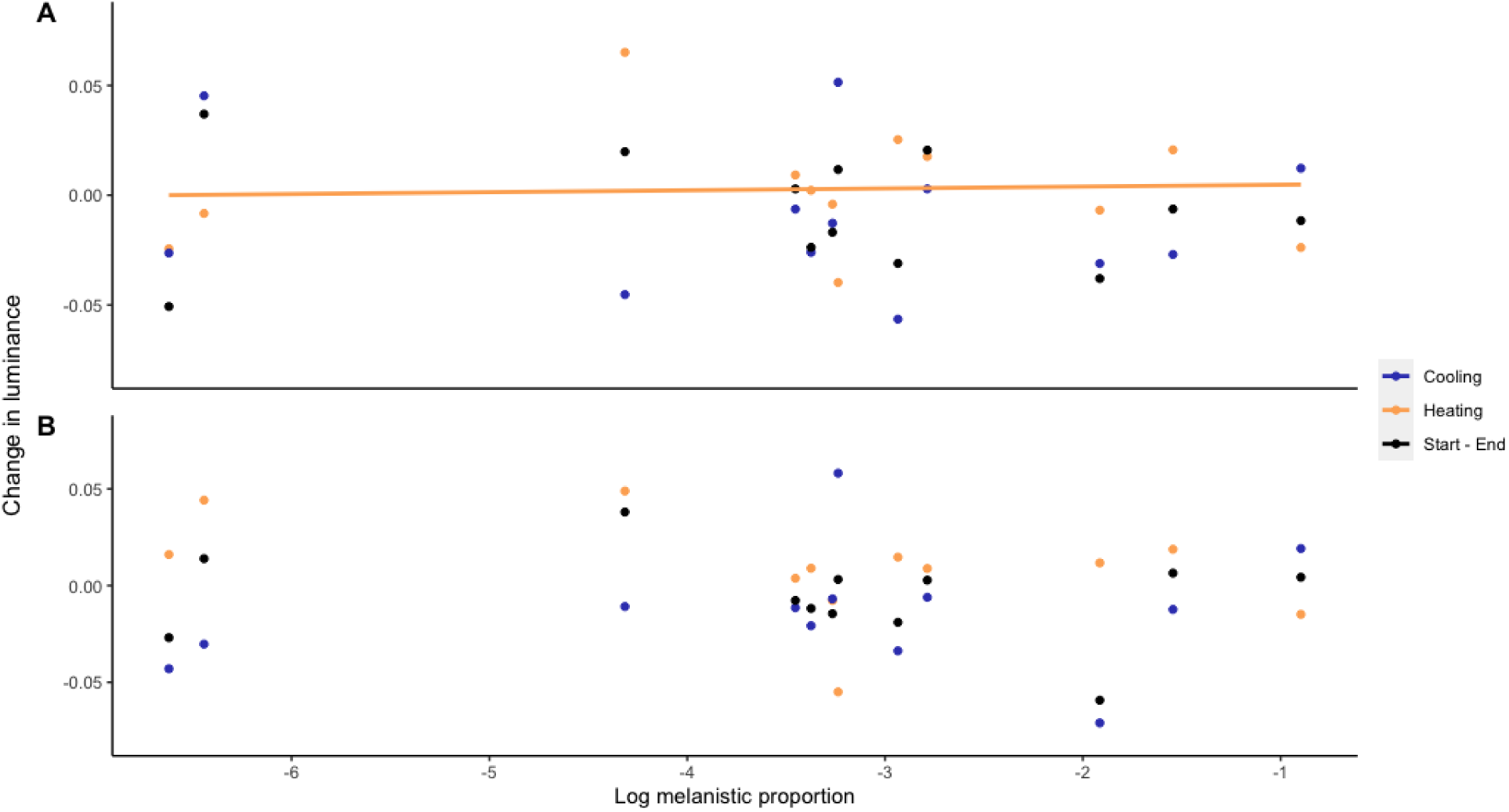
Change in average luminance for the non-melanistic (A) and melanistic (B) part of the body against melanistic proportion plotted on a logarithmic scale. Average luminanc was calculated by weighted RGB values taken from images. Heating rates were calculated by taking the average body temperature difference between each block for blocks 5 and 3 per individual. Cooling rates were calculated using the same methods for blocks 1 and 3. Each dot represents an individual. Blue dots refer to the cooling phase (blocks 1-3), orange dots to the heating phase (blocks 3-5), and black dots refer to the start and end of the experiment (blocks 1- 5). Solid line indicates a significant p-value via a linear model.

**Table 4.** Influence of melanistic proportion and body temperature on changes in dorsal luminance during cooling (blocks 1-3), heating (blocks 3-5), and beginning and end of the experiment (blocks 1-5) for the melanistic (a) and non-melanistic (b) dorsal areas of the body. Melanistic proportion was calculated as the area of stable melanistic color relative to the entire dorsal area, averaged across all images of a single individual and transformed to a log scale for analyses. Significant p-values (<0.05) are indicated in bold. * indicates interaction of variables tested in the models.

| Blocks | covariates | Estimate | Std Err | t-value | p-value |
| --- | --- | --- | --- | --- | --- |
| a) Change in luminance for the melanistic area of the body |  |  |  |  |  |
| 1-3 | Temperature | 0.0215 | 0.0293 | 0.733 | 0.49 |
|  | Melanistic proportion | 0.1826 | 3.1282 | 0.058 | 0.96 |
|  | Temperature*melanistic proportion | 0.0077 | 0.3685 | 0.021 | 0.98 |
| 3-5 | Temperature | -0.0348 | 0.0180 | -1.934 | 0.09 |
|  | Melanistic proportion | -1.1748 | 0.7829 | -1.501 | 0.17 |
|  | Temperature*melanistic proportion | 0.1683 | 0.1157 | 1.455 | 0.18 |
| 1-5 | Temperature | -0.0108 | 0.01429 | -0.755 | 0.47 |
|  | Melanistic proportion | 0.3099 | 0.1655 | 1.872 | 0.10 |
|  | Temperature*melanistic proportion | 0.1936 | 0.1046 | 1.851 | 0.10 |
| b) Change in luminance for the non-melanistic area of the body |  |  |  |  |  |
| 1-3 | Temperature | -0.0348 | 0.0353 | -0.984 | 0.35 |
|  | Melanistic proportion | -0.1337 | 0.0860 | -1.554 | 0.16 |
|  | Temperature*melanistic proportion | -0.0168 | 0.0109 | -1.547 | 0.16 |
| 3-5 | Temperature | 0.0293 | 0.0168 | 1.740 | 0.12 |
|  | Melanistic proportion | -0.1164 | 0.0407 | -2.861 | **0.02** |
|  | Temperature*melanistic proportion | 0.0177 | 0.0062 | 2.877 | **0.02** |
| 1-5 | Temperature | 0.0077 | 0.0216 | 0.357 | 0.73 |
|  | Melanistic proportion | -0.0020 | 0.0117 | -0.176 | 0.87 |
|  | Temperature*melanistic proportion | -0.0019 | 0.0086 | -0.221 | 0.83 |

## DISCUSSION

Crepuscular and nocturnal reptiles experience low exposure to solar radiation during their active times and have been suggested to primarily rely on thigmothermy. Tigmothermy refers to the absorption of heat from the surrounding environment, rather than directly from solar radiation (Garrick, 2008; M. Kearney & Predavec, 2000). Despite relying mostly on substrate and surrounding temperatures, crepuscular and nocturnal reptiles can also bask for thermoregulation when needed (Angilletta et al., 1999).

In this study, we directly compare how substrate and atmospheric temperatures correlate with the internal body temperature in *E. macularius* and the role of melanistic pattern on physiological color change and thermoregulation in these animals. In our study, the main sources of heat during the cooling down of our experiments (blocks 1-3) were provided by the heat pad placed at one side of the terrarium and the lamp placed above the terrarium. We found a stronger correlation between body temperatures and terrarium substrate temperatures than for atmospheric temperatures, suggesting that *E. macularius* relies on absorbing heat from the ground and is thigmothermic. To notice that while the substrate temperature of the spot at which the gecko was located and the body temperatures were both estimated from IR photographs, the median atmospheric temperature - as estimated from data loggers - was generally lower than the gecko body temperature and the substrate temperature estimated from data loggers, suggesting that the higher correlation observed between substrate temperatures and body temperatures is not due to how the temperatures were measured. Our findings also support what previously suggested (Craioveanu et al., 2017; Garrick, 2008) based on measures of body surface temperature and ambient temperature differentials.

Although *E. macularius* mostly rely on the substrate temperature for thermoregulation, melanistic coloration – both as a stable coloration and as a physiological darkening of the skin – can be used for thermoregulation to increase or decrease the animal’s body temperature during basking (Angilletta et al., 1999; Clusella Trullas et al., 2007; Smith, Cadena, Endler, Kearney, et al., 2016; Smith, Cadena, Endler, Porter, et al., 2016). Specifically, melanin can be used to absorb UV radiation and convert it into heat by photon-phonon transformation (McNamara et al., 2021) thus contributing to increased heating rates, but can also increase the rate of heat transfer for cooling in reptiles (Geen & Johnston, 2014). as most research into melanism has been conducted Compared to what we know on the role of melanistic coloration on diurnal ectotherms (Belliure & Carrascal, 2002; Garrick, 2008), the function(s) of melanistic coloration and especially of melanistic pattern in crepuscular and nocturnal reptiles are largely unknown. Our results based on 12 individuals of *E. macularius* with different proportions of melanistic pattern, indicate that melanistic proportion is independent from body size (SVL), body weight, or sex, and that body size (SVL), body weight, or sex do not influence the heating or cooling rates. We found that the proportion of melanistic pattern does not influences cooling or heating rates in *E. macularius*. As *E. macularius* is crepuscular and only active for a few hours of daylight and mostly relies on thigmothermy, the characteristic spotted melanistic pattern of *E. macularius* in nature may be used for purposes different from thermoregulation.

However, exposure to prolonged suboptimal low temperatures, as tested here, may trigger a physiological response to darken the skin color to increase heat absorbance. This phenomenon to date has only been observed in heliothermic reptiles – reptiles that regulate body temperature through solar radiation (Cowles, 1940). As *E. macularius* is crepuscular and shelters during daylight hours (Angilletta et al., 1999), physiological color change may be used more for cooling purposes, for example to increase heat transfer to the environment if shelters exceed optimal temperatures during daylight. In our study, we found that luminance (changes in luminance corresponds to physiological color change) does not change significantly between blocks during cooling temperatures, but it does during the heating phases of the experiment for the non-melanistic part of the body and depending on the melanistic proportion of the animals. We found that for the non-melanistic areas of the body, individuals with higher proportions of melanistic pattern experienced less darkening of the skin, and vice-versa. Physiological color changes during heating – along with other mechanisms such as changes in peripheral blood flow (Bartholomew et al., 1965; Rice & Bradshaw, 1980) - may provide protection against overheating, as the rate of heat gain has been observed to increase at higher environmental temperatures (Belliure & Carrascal, 2002). A caveat of these results is however that the significant relationship between melanistic proportion, heating, and physiological color change are strongly influenced by one individual with high melanistic proportion.

Taken together, our results suggest that in *E. macularius* melanistic pattern may not be used for thermoregulation, while physiological color change may occur to prevent overheating. Further research is needed to understand the role of melanistic coloration and melanistic pattern in geckos and the extent to which physiological color change occurs, especially in crepuscular and nocturnal species. In geckos, spotted patterns – such as the one observed in *E. macularius* – have been suggested to represent a more specialized type of camouflage, although bands and not spots have been proposed to be associated with nocturnal activity (Allen et al., 2020). Previous studies on crepuscular and nocturnal geckos have also indicated that physiological color change may be used more for background matching and camouflage than for thermoregulation (Vroonen et al., 2012; Zaidan III & Wiebusch, 2007). The few studies investigating physiological color change for thermoregulation and camouflage in other nocturnal geckos propose that these functions are mutually exclusive phenomena (Vroonen et al., 2012; Zaidan III & Wiebusch, 2007), but based on our results, this may dependent on the occurrence and proportion of melanistic pattern. Future studies should therefore investigate the functional trade-off between melanistic coloration, including melanistic pattern and physiological color change, for thermoregulation versus its use in camouflage or signaling in crepuscular and nocturnal reptiles in captivity and in the wild. Finally, although our study is based on captive bread animals and wild *E. macularius* do not show the same extent of variation in melanistic proportion as for the captive bred animals, our study is noteworthy mostly from a methodological and theoretical point of view. First, we developed a freely available software package that can be used to extract color patterns information from digital images of freely moving organisms with soft bodies, as in the case of geckos. Secondly, our results highlight the importance of melanistic pattern for study on thermoregulation and coloration and suggest that melanistic pattern and melanistic coloration may be used for multiple non-exclusive functions. The conclusions of our study could help with further understanding the function of coloration and color pattern development in nocturnal and crepuscular reptiles, and how their function may differ from those of diurnal reptiles.

## Acknowledgments

We are thankful to Emanuele Scanarini, Tony Gamble, Andran Abramjan, Alyssa Stark, Scott Glaberman, and Pat Gillevet for helping with the experimental setup. Andrea Weeks and Daniel Hanley provided helpful comments on an earlier version of this paper.

## Competing interests

The authors declare no competing interests

## Funding

George Mason University, Office of Student Scholarship, Creative Activities, and Research (OSCAR).

## Data availability

Full dataset will be available on Dryad after manuscript acceptance.

## Code availability

https://github.com/brandon-hastings/Lumeleon

## Notes

### Competing Interest Statement

The authors have declared no competing interest.

